# Pathogen evasion of chemokine response through suppression of CXCL10

**DOI:** 10.1101/557876

**Authors:** Alejandro L. Antonia, Kyle D. Gibbs, Esme Trahair, Kelly J. Pittman, Benjamin H. Schott, Jeffrey S. Smith, Sudarshan Rajagopal, J. Will Thompson, R. Lee Reinhardt, Dennis C. Ko

**Author notes:** To whom correspondence should be addressed: Dennis C. Ko, 0048B CARL Building Box 3053, 213 Research Drive, Durham, NC 27710. @denniskoHiHOST.

## Abstract

Clearance of intracellular pathogens, such as *Leishmania* (*L.*) *major*, depends on an immune response with well-regulated cytokine signaling. Here we describe a pathogen-mediated mechanism of evading CXCL10, a chemokine with diverse antimicrobial functions, including T cell recruitment. Infection with *L. major* in a human monocyte cell line induced robust *CXCL10* transcription without increasing extracellular CXCL10 protein concentrations. We found that this transcriptionally independent suppression of CXCL10 is mediated by the virulence factor and protease, glycoprotein-63 (*gp63)*. Specifically, GP63 cleaves CXCL10 after amino acid A81 at the base of a C-terminal alpha-helix. Cytokine cleavage by GP63 demonstrated specificity, as GP63 cleaved CXCL10 and its homologues, which all bind the CXCR3 receptor, but not distantly related chemokines, such as CXCL8 and CCL22. Further characterization demonstrated that CXCL10 cleavage activity by GP63 was produced by both extracellular promastigotes and intracellular amastigotes. Crucially, CXCL10 cleavage impaired T cell chemotaxis *in vitro*, indicating that cleaved CXCL10 cannot signal through CXCR3. Ultimately, we propose CXCL10 suppression is a convergent mechanism of immune evasion, as *Salmonella enterica* and *Chlamydia trachomatis* also suppress CXCL10. This commonality suggests that counteracting CXCL10 suppression may provide a generalizable therapeutic strategy against intracellular pathogens.

**Importance:** Leishmaniasis, an infectious disease that annually affects over one million people, is caused by intracellular parasites that have evolved to evade the host’s attempts to eliminate the parasite. Cutaneous leishmaniasis results in disfiguring skin lesions if the host immune system does not appropriately respond to infection. A family of molecules called chemokines coordinate recruitment of the immune cells required to eliminate infection. Here, we demonstrate a novel mechanism that *Leishmania (L.) major* employs to suppress host chemokines: an *L. major* protease cleaves chemokines known to recruit T cells that fight off infection. We observe that other common human intracellular pathogens, including *Chlamydia trachomatis* and *Salmonella enterica*, reduce levels of the same chemokines, suggesting a strong selective pressure to avoid this component of the immune response. Our study provides new insights into how intracellular pathogens interact with the host immune response to enhance pathogen survival.

## Introduction

Proper immune clearance of intracellular pathogens requires precise cytokine and chemokine signaling. These cytokines coordinate the localization, activation, and polarization of innate and adaptive immune cell subsets. To study T cell recruitment and polarization in response to intracellular pathogens, parasites in the genus *Leishmania* have served as a paradigm (1). However, persistent gaps in the understanding of host and pathogen factors that influence T cell response and recruitment contribute to the dearth of immunotherapeutics and vaccines. With no available vaccine and limited treatment options, *Leishmania* spp. continue to cause 1.2 million cases of cutaneous leishmaniasis (CL) and 0.4 million cases of visceral leishmaniasis annually (VL) (2). A better understanding of host immunity and pathogen evasion strategies is imperative to develop alternative approaches to current therapies, which are limited by variable efficacy, high cost, and growing drug resistance (3). Of particular relevance may be instances where multiple diverse pathogens have evolved to evade or suppress the same key host immune signaling pathways (4, 5).

To clear *L. major* parasites, a causative agent of CL, the adaptive immune system must be coordinated to a type-1 response by appropriately recruiting immune cell subsets, particularly CD4+ T helper 1 (T_h_1) cells and CD8+ cytotoxic T lymphocytes (CTLs) (6). This recruitment is mediated by chemokines, a family of signaling molecules that regulate recruitment and localization of unique immune cell subsets. For example, T_h_1 cells, which mediate a pro-inflammatory response effective at eliminating intracellular parasites, are recruited by chemokines such as CXCL10 through the CXCR3 receptor. By contrast, T_h_2 cells, which promote immunity targeting extracellular parasites, are recruited by chemokines such as CCL22 through the CCR4 receptor (7). When infected with *L. major*, T_h_2 responding mice develop non-healing lesions, whereas T_h_1 responding mice effectively clear the parasite (8–10). As part of the broader type-1 immune response against *L. major* infection, parasite-specific CD8+ cells are also recruited, and have been implicated in productive immunity to primary and secondary infection (11–14). Corresponding observational studies in humans support this model where non-healing cutaneous lesions are characterized by T_h_2 associated cytokines, and individuals resistant to lesion development have a higher predominance of T_h_1 associated cytokines (15–18). Together these studies highlight the critical role of cytokine and chemokine signaling in specific immune cell subsets during infection.

One of the chemokines that specifically regulates localization and activity of CD4+ T_h_1 and effector CD8+ T-cells is CXCL10, or IFN_γ_ Inducible Protein 10 (IP10). CXCL10 is part of a family of highly homologous chemokines, including CXCL9 and CXCL11, which bind to and activate the CXCR3 chemokine receptor (reviewed in (19)). Multiple lines of investigation suggest that CXCL10 protects against *Leishmania* infection. First, the host upregulates *CXCL10* transcription throughout infection (20–22) and cells expressing CXCR3 are expanded after infection (23). Second, BALB/c mice, which are unable to control *Leishmania* spp. infection, demonstrate a defect in CXCR3 upregulation (24, 25). Finally, exogenous CXCL10 is protective against both cutaneous and visceral leishmaniasis (26–29). Therefore, the type-1 associated chemokine CXCL10 is important for host control of cutaneous leishmaniasis.

Beyond *Leishmania*, type-1 immunity and CXCL10-CXCR3 signaling are critical for clearing other intracellular pathogens. For the obligate intracellular bacterium *Chlamydia trachomatis*, T_h_1 cells are required for clearance of infection while a T_h_2 dominated response may lead to excessive pathology (30–33). In mice, CXCL10 mRNA and protein are significantly induced after infection (34–36). Similarly, T_h_1 responses are crucial for an effective immune response to the facultative intracellular bacteria *Salmonella enterica* serovar Typhimurium based on studies in mice (37, 38) and the predisposition of people with rare mutations in T_h_1-promoting cytokines (IFN_γ_ and IL12) to invasive Salmonellosis *(39).* Further, M1-polarized macrophages, which restrict *Salmonella* intracellular replication (40, 41), robustly upregulate *CXCL10* transcription (42, 43). Finally, mice deficient for CXCR3 have increased susceptibility to *S. enterica (44)*, *Toxoplasma* (*T.*) *gondii* (45), and *C. trachomatis* (46). Thus, the CXCL10-CXCR3 signaling axis coordinates an adaptive type-1 immune response to intracellular pathogens that promotes a successful healing response.

Here, we report that *L. major* suppresses extracellular CXCL10 protein levels, providing a potential mechanism for evasion of the adaptive immune response. This suppression occurs through the proteolytic activity of the virulence factor glycoprotein-63 (GP63). GP63 cleavage of CXCL10 occurs throughout *in vitro* infection and abrogates CXCR3-dependent T cell migration. Furthermore, we observed CXCL10 suppression with other intracellular pathogens, including *S. enterica* and *C. trachomatis*, demonstrating that diverse intracellular pathogens have developed convergent mechanisms to suppress CXCL10.

## Results

### *L. major* infection suppresses CXCL10 protein, despite induction of *CXCL10* mRNA

To broadly screen for *L. major* manipulation of host immunity, we measured secreted levels of 41 cytokines following infection of lymphoblastoid cell lines (LCLs) with *L. major.* LCLs constitutively produce CXCL10, and incubation with *L. major* reduced CXCL10 levels by greater than 90% (Fig. 1A). We confirmed this decrease in CXCL10 protein in LPS-stimulated human THP-1 monocytes infected with *L. major* (Fig. 1B). Despite the reduction in CXCL10 protein in culture supernatants, THP-1s exposed to *L. major* had 2.5-fold higher *CXCL10* mRNA relative to uninfected (Fig. 1B). Therefore, *L. major* suppresses CXCL10 protein through a transcriptionally independent mechanism.

**Figure 1.**
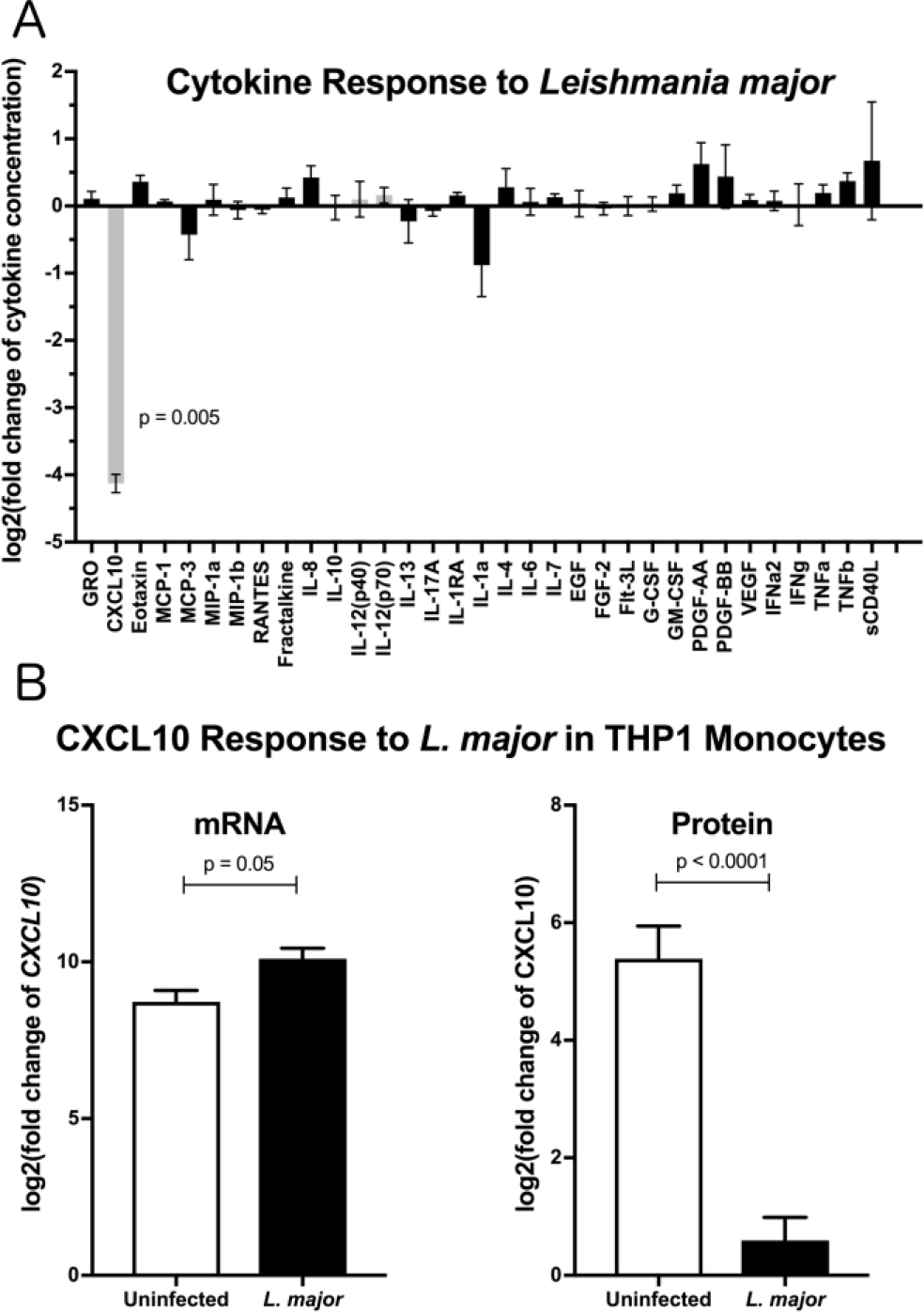
*Leishmania major* suppresses CXCL10 post-transcriptionally in multiple human cell lines. (A) Cytokine screening of LCLs exposed to *L. major* demonstrated suppression of CXCL10. Three lymphoblastoid cell lines (LCL), 7357, 18524, and 19203, were infected with *L. major.* Chemokines secreted into culture supernatants were analyzed by Luminex. Cytokines below the limit of detection were removed from the final analysis. Values are represented as log_2_ of the fold change relative to uninfected LCLs. Type-1 associated cytokines are represented in grey. P value represents Dunnett’s post-hoc test compared to 1, after repeated measures one-way ANOVA. (B) CXCL10 produced by LPS stimulated THP-1 monocytes was suppressed by *L. major.* THP-1 monocytes were stimulated with LPS prior to *L. major* infection. CXCL10 mRNA was measured by qRT-PCR TaqMan assay using the ΔΔC_t_ method with 18s as housekeeping gene, and CXCL10 protein was measured by ELISA. For mRNA (n=3) and ELISA (n=6), Fold Change is relative to unstimulated, uninfected THP-1s. P values calculated by Student’s *t-test*.

### The *L. major* matrix-metalloprotease, glycoprotein-63 (GP63), is necessary and sufficient for CXCL10 protein suppression

To test whether an *L. major*-secreted factor is responsible for CXCL10 protein suppression, we treated recombinant human CXCL10 with cell-free conditioned media obtained from cultured *L. major* promastigotes. Again, CXCL10 was reduced by 90% with the conditioned media (Fig. 2A). These results were consistent with proteolytic degradation by a pathogen-secreted protease. We hypothesized that CXCL10 suppression was mediated by glycoprotein-63 (GP63), a zinc-metalloprotease conserved among the *Trypanasoma* family of parasites and expressed in both the extracellular promastigote and intracellular amastigote life stages (47–50). To test if GP63 is required to suppress CXCL10, we used a known GP63 inhibitor, the zinc-chelator 1, 10-phenanthroline (51). 1,10-phenanthroline inhibited CXCL10-suppressive activity in *L. major* conditioned media (Fig. 2A). Consistent with GP63-mediated degradation of CXCL10, conditioned media from a promastigote culture of *L. major* deficient for *gp63* (Δ*gp63;* (52)) did not suppress CXCL10, whereas complementation with a single copy of *gp63* (*L. major* Δ*gp63*+1) restored CXCL10 suppression (Fig. 2B). Furthermore, heterologously expressed GP63 secreted from mammalian HEK293T cells was sufficient for complete CXCL10 suppression, while a single point mutation in the catalytic site of GP63 (E265A) abrogated suppression (Fig. 2C). Therefore, GP63 is both necessary and sufficient for CXCL10 suppression by *L. major.*

**Figure 2.**
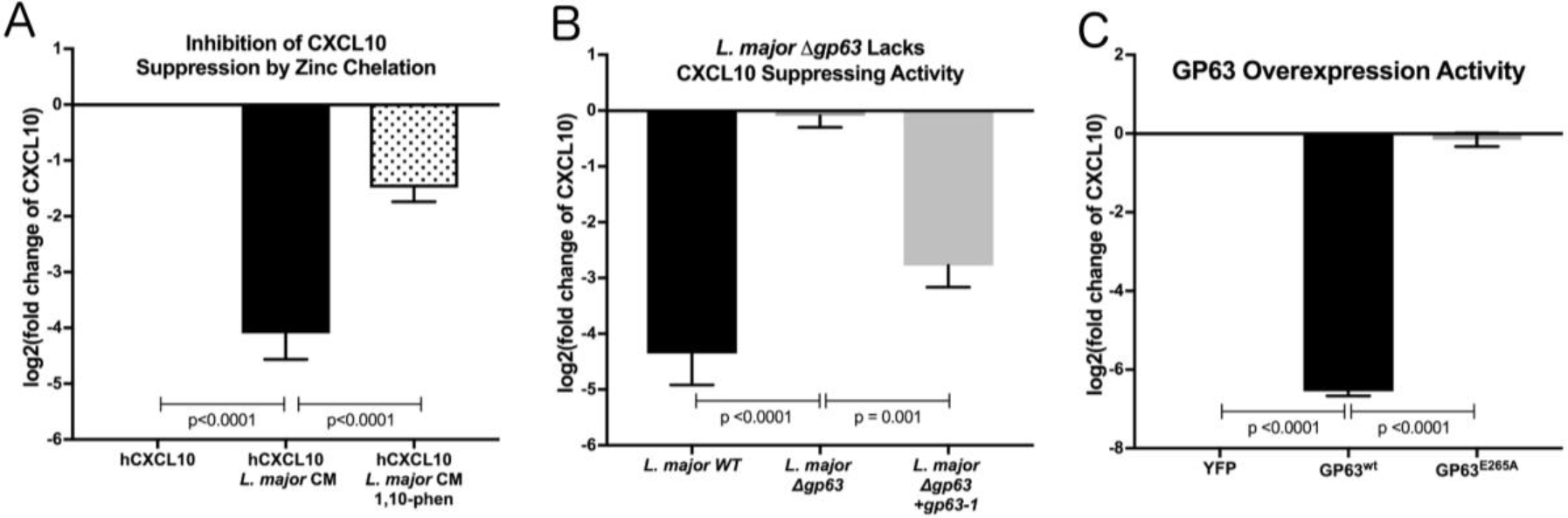
*Leishmania major* matrix-metalloprotease, glycoprotein-63, is necessary and sufficient to cleave CXCL10. (A) Zinc chelation prevents CXCL10 suppression. Concentration of human recombinant CXCL10 was measured by ELISA after incubation for 12 hours with filtered conditioned media from *L. major* WT promastigote culture and addition of the zinc-chelator 1,10-phenanthroline (n=8). (B) *gp63* is required for *L. major* CXCL10 suppression. Human recombinant CXCL10 concentrations were measured by ELISA after 12 hour incubation with conditioned media from *L. major* WT, Δ*gp63*, or Δ*gp63*+*1* (n=6). (C) GP63 expressed and secreted by HEK293Ts is sufficient for CXCL10 suppression. Human recombinant CXCL10 concentrations were measured by ELISA after 12 hour incubation with culture supernatant from HEK293Ts transfected with pCDNA3.1-gp63^WT^ or pCDNA3.1-gp63^E285A^ (n=7). P values calculated by one-way ANOVA with Tukey’s post-hoc test. Error bars represent standard error of the mean.

### GP63 selectively cleaves the CXCL10-related family of chemokines at the start of the C-terminal alpha-helix

As GP63 has a diverse set of identified *in vitro* substrates (48), we examined the specificity of GP63 across a spectrum of chemokines. Based on the initial cytokine screen (Figure 1A), we hypothesized GP63 cleavage would be restricted to CXCL10 and highly related chemokines. To experimentally test for GP63 cleavage, purified recombinant chemokines were incubated with conditioned media from *L. major* WT, *L. major* Δ*gp63*, or *L. major* Δ*gp63*+*1*. GP63 cleavage was observed for CXCL9 (38.14% amino acid identity with CXCL10) and CXCL11 (30.85% amino acid identity with CXCL10) (Figure 3A, B), which both signal through CXCR3. By contrast, no cleavage of CXCL8 (IL-8; a neutrophil-attracting chemokine) or CCL22 (MDC; a T_h_2-attracting chemokine) was detected (Fig. 3B). Thus, chemokine cleavage by GP63 appears to preferentially degrade chemokines involved in CXCR3 signaling.

**Figure 3.**
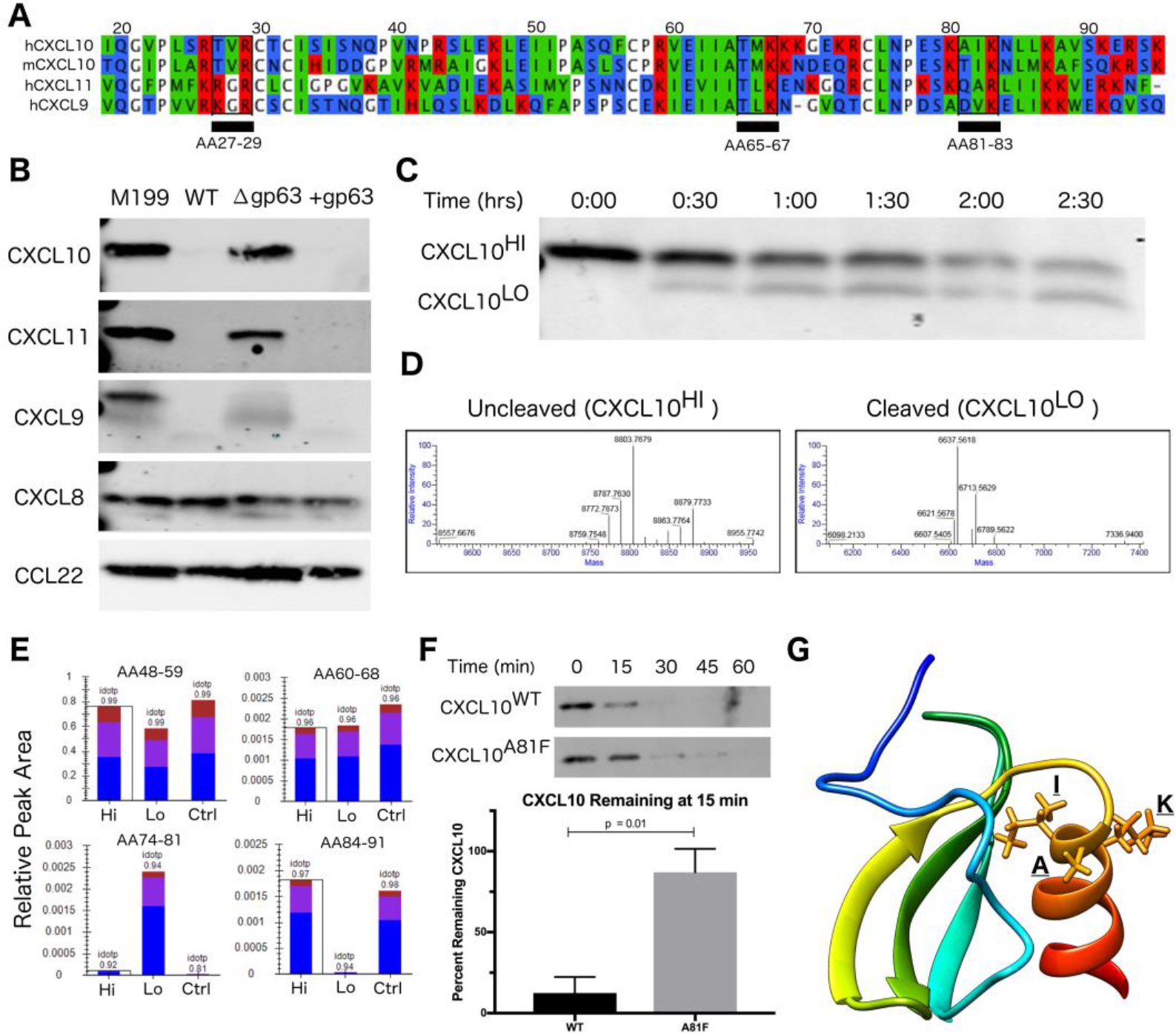
CXCL10 cleavage by GP63 occurs between positions A81 and I82. (A) CXCL9/10/11 share significant homology at the amino-acid level. Multisequence alignment demonstrates that physical characteristics of amino acids are conserved across the CXCL10 family of chemokines. There are three putative GP63 cleavage sites (underlined) based on the consensus sequence of polar (P1), hydrophobic (P1’), basic (P2’) (53). (B) GP63 selectively cleaves chemokine ligands of the CXCR3 receptor. Conditioned media from *L. major WT, Δgp63*, and *Δgp63*+*1* was incubated with human recombinant chemokines for 12 hours and product detected by western blot. Cleavage is only detected for the CXCL9/10/11 family. (C) Cleavage by GP63 generates a smaller molecular weight protein. A time course of cleavage of human CXCL10 by heterologously expressed GP63 demonstrated an intermediate cleavage product, resolved by PAGE and Coomassie staining. (D) Cleavage by GP63 results in a change in CXCL10 molecular weight of 2.2kD. Capillary electrophoresis-Mass Spectrometry (CE-MS) determined the molecular weight of the uncleaved (CXCL10^Hi^) and cleaved (CXCL10^Lo^) bands as 8.8kD and 6.6kD respectively. (E) Comparative analysis by trypsin digest of cleaved and uncleaved CXCL10 reveals cleavage occurring between A81-I82. Liquid chromatography-mass spectrometry (LC-MS) following trypsin digest of CXCL10^Hi^ and CXCL10^Lo^ identified peptide ending at A81, exclusively in the CXCL10^Lo^ band, and a corresponding lack of peptide coverage from AA84-91. (F) Mutation of A81F significantly impairs GP63 cleavage of CXCL10. In the presence of GP63, CXCL10^A81F^ remains stable for up to 45 minutes whereas CXCL10^WT^ degradation is nearly complete by 15 minutes. Percentage of GP63 remaining at 15 minutes is plotted (n=3-4 per CXCL10 genotype). P value calculated by Student’s *t-test*. (G) The GP63 cleavage site is found on the C-terminal alpha-helix loop of CXCL10. Based on the NMR crystal structure of CXCL10 (Booth et al., 2002), the A81, I82, K83 (P1, P1’, P2’) GP63 cleavage motif maps to an exposed alpha-helical region.

Although western blot analysis supported GP63-dependent cleavage through loss of CXCL10 immunoreactivity, it did not reveal the cleavage site or potential cleavage products. The GP63 consensus cleavage site has been described as polar, hydrophobic, and basic amino acids at positions P1, P1’, and P2’ (53). Following this pattern, there are three potential cleavage sites in the mature CXCL10 protein (from amino acid position 22-96) that are conserved between human and murine CXCL10 (68.37% amino acid identity) (Fig. 3A). In order to characterize the cleavage product(s), we incubated GP63 with human recombinant CXCL10 and visualized a shift in size by total protein stain (Fig. 3C). Intact CXCL10 and the largest cleavage products were determined to be 8.8kD and 6.6kD respectively, by capillary electrophoresis-mass spectrometry (CE-MS) (Fig. 3D). After running the sample on a PAGE gel, the 8.8kD (intact, “Hi”) product, 6.6kD (cleaved, “Lo”) product, and an uncleaved control (“Ctrl”) were sequenced by trypsin digestion followed by liquid chromatography-tandem mass spectrometry (LC-MS/MS). Comparison of peptides after trypsin digest revealed a peptide from amino acids (AA) 74-81 that was exclusively present in the cleaved CXCL10 band, but notably absent in the uncleaved band (Fig. 3E). Conversely, distal peptide fragments such as AA84-91 were only present in the uncleaved CXCL10. This analysis demonstrated cleavage occurring in between A81 and I82, resulting in the loss of detectable peptides beyond those amino acids in cleaved CXCL10. This is consistent with the fragment size based on intact molecular weight CE-MS, and notably AIK (AA 81-83) is one of the three potential cleavage sites identified in our comparative analysis (see Figure 3A). To confirm this site as preferred for GP63 cleavage, we used site-direct mutagenesis to mutate the residues in the proposed cleavage motif. Mutation of the identified P1 residue significantly slowed CXCL10 cleavage in a time course experiment (Fig. 3F). Mapping the residues onto the crystal structure of CXCL10 (54) demonstrated that cleavage occurs at the beginning of the C-terminal alpha-helix of CXCL10 (Fig. 3G).

### GP63 produced by *L. major* promastigotes or amastigotes can cleave CXCL10 protein

Immediately after injection by the sand-fly vector, *Leishmania* parasites exist as an extracellular, flagellated promastigote but are rapidly phagocytized where they transform into the intracellular, aflagellated amastigote parasite stage. We hypothesized that GP63 would continue to be able to suppress CXCL10 through both stages of infection, as transcriptomics indicate GP63 expression during both stages (50). To test the capacity of *L. major* to suppress CXCL10 in both the promastigote and amastigote stage of infection, we utilized PMA differentiated THP-1 monocytes as an intracellular macrophage model of infection. Differentiated THP-1 monocytes were infected at an MOI of 20 with promastigotes from *L. major WT, Δgp63*, or *Δgp63*+*1*. Extracellular promastigote activity was assessed in the supernatant at 24 hours post infection, followed immediately by washing to remove the extracellular promastigotes and GP63 protein in the media, and subsequently assessing intracellular amastigote activity at 48 hours post infection.

This model demonstrated CXCL10 protein suppression by GP63 in both stages of the parasite life cycle. *L. major WT* promastigotes had no induction of CXCL10 protein relative to uninfected cells, while *L. major Δgp63* infection resulted in a significant induction of CXCL10 protein (Fig. 4A). Similarly, the *L. major WT* amastigotes continued to suppress CXCL10 protein while *L. major Δgp63* infection significantly induced CXCL10 protein (Fig. 4B). The complementation observed with the *L. major Δgp63*+*1* strain is significant, though incomplete in the promastigote stage and further reduced in the amastigote stage. This is attributable to the plasmid construct being designed for high expression in the promastigote stage (55) and the lack of G418 selection during the experiment. Notably, all three *L. major* strains cause comparable induction of *CXCL10* mRNA (Fig. 4C). These results indicate that CXCL10 mRNA is induced during *Leishmania* infection, but protein levels are reduced by GP63, present at both parasite life cycle stages involved in infection in mammalian hosts.

**Figure 4.**
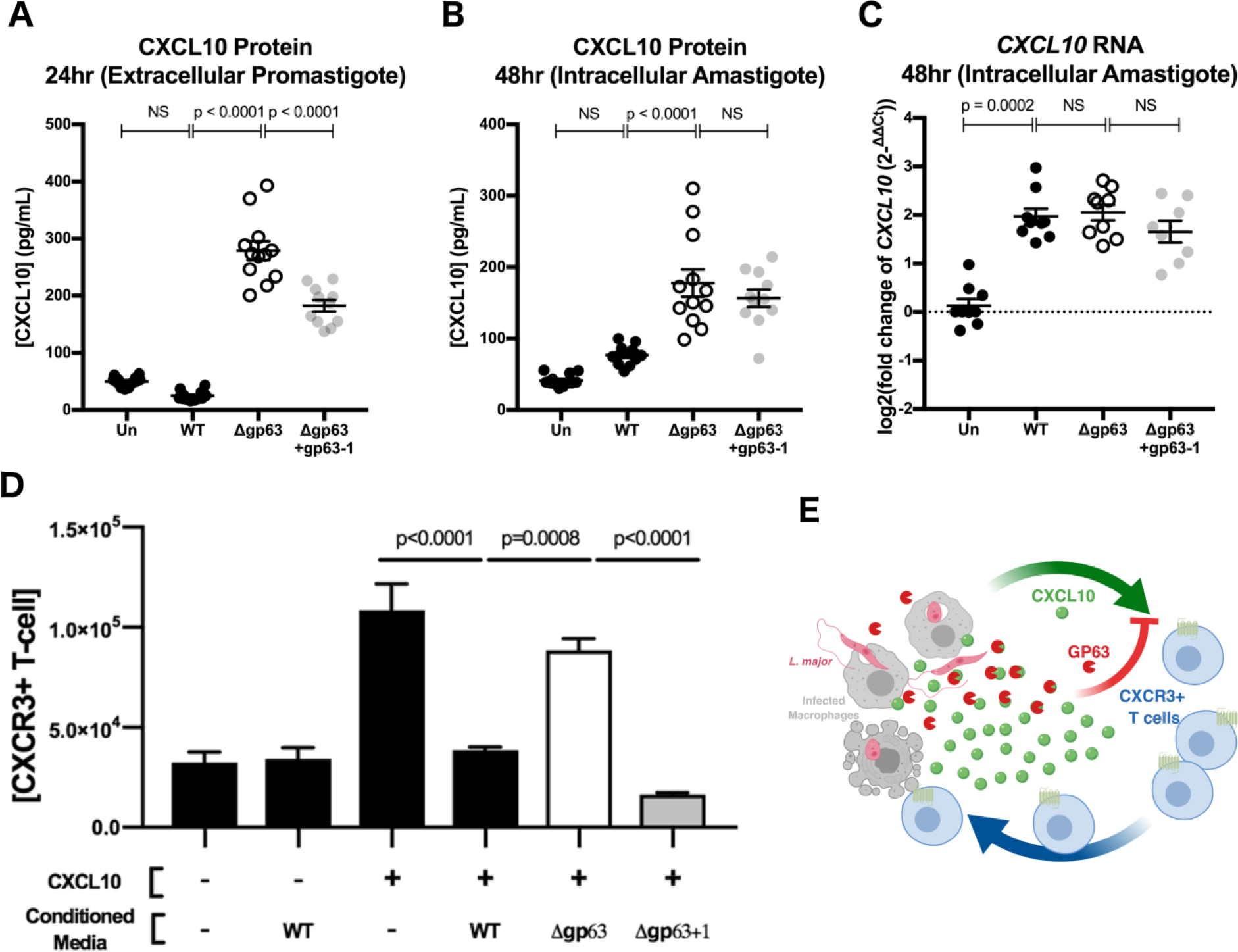
GP63 produced by *L. major* promastigotes and amastigotes cleaves CXCL10 and abolishes its chemotactic activity. (A) *L. major* promastigotes suppress CXCL10 through GP63 activity. THP-1 monocytes were differentiated using 100ng/mL of PMA prior to infection and CXCL10 concentration was assessed in the supernatant 24 hours post-infection by ELISA. (B) *L. major* amastigotes suppress CXCL10 through GP63 activity. At 24 hours post-infection, extracellular promastigotes were washed away from the differentiated THP-1 monocytes. At 48 hours post-infection the CXCL10 concentration was assessed in the supernatant by ELISA. For A-B, data represents four separate infections and was analyzed by one-way ANOVA with Tukey’s post-hoc test (C) *L. major* induces similar levels of *CXCL10* mRNA, independent of GP63 genotype. At 48 hours post-infection, mRNA was extracted from PMA differentiated THP-1 monocytes and CXCL10 mRNA was measured by qRT-PCR TaqMan assay using the ΔΔC_t_ method with 18s as housekeeping gene. For C, data are from three separate experiments and were analyzed by one-way ANOVA with Tukey’s post-hoc test. (D) CXCL10 incubated with GP63 is unable to chemoattract CXCR3+ cells *in vitro.* Jurkat T cells stably transfected with CXCR3 were seeded on the apical surface of a 5μm transwell insert, with human recombinant CXCL10 pre-incubated with conditioned media from either *L. major WT, Δgp63*, or *Δgp63*+*1* in the basal chamber. The number of CXCR3+ Jurkats in the basal chamber after 4 hours were counted to assess chemotactic capacity of CXCL10 after exposure to GP63. (E) Proposed model where the host attempts to upregulate CXCL10 in response to infection, but through the activity of GP63 *L. major* is able to impair signaling through the CXCR3 receptor.

### GP63 cleaved CXCL10 is unable to recruit CXCR3 expressing T cells

Because CXCL10 coordinates the recruitment of CXCR3+ T cells during infection, we next tested if GP63 cleavage of CXCL10 impacts T cell recruitment. We tested the chemotactic ability of cleaved CXCL10 to recruit Jurkat T cells expressing CXCR3. The basal chamber of a transwell system was seeded with CXCL10 in the presence of conditioned media from *L. major WT, L. major Δgp63*, or *L. major Δgp63*+*1*. Conditioned media from *L. major* WT and *L. major Δgp63*+*1* abrogated CXCL10 induced migration of CXCR3+ Jurkat T cells, whereas the *L. major Δgp63* conditioned media did not impair chemotaxis (Fig. 4D). Together these data support a model whereby the host attempts to produce CXCL10 to coordinate recruitment of CXCR3 expressing immune cells, but *L. major* produces GP63 to inactivate CXCL10 and impair T cell chemotaxis (Fig. 4E).

### CXCL10 suppression has evolved independently in multiple intracellular pathogens

Given that CXCL10 mediates a type-1 immune response that protects against a broad range of intracellular pathogens, we asked if suppression of CXCL10 has evolved in has evolved in other parasites and bacteria. CXCL10 production by LCLs was measured by ELISA after exposure to a variety of pathogens including *Toxoplasma* (*T.*) *gondii*, *Plasmodium* (*P.*) *bergei*, *Salmonella* (*S.*) *enterica* serovar Typhimurium, *Chlamydia* (*C.*) *trachomatis*, *Mycobacterium* (*M.*) *marinum, Mycobacterium* (*M.*) *smegmatis*, *Staphylococcus* (*S.*) *aureus*, and *Cryptococcus* (*C.*) *neoformans.* CXCL10 suppression of at least 80% was observed with two additional intracellular pathogens: *S.* Typhimurium and *C. trachomatis.* In contrast, other pathogens, including the extracellular pathogens *S. aureus* and *C. neoformans*, exhibited modest to no suppression of CXCL10 (Fig. 5A).

**Figure 5.**
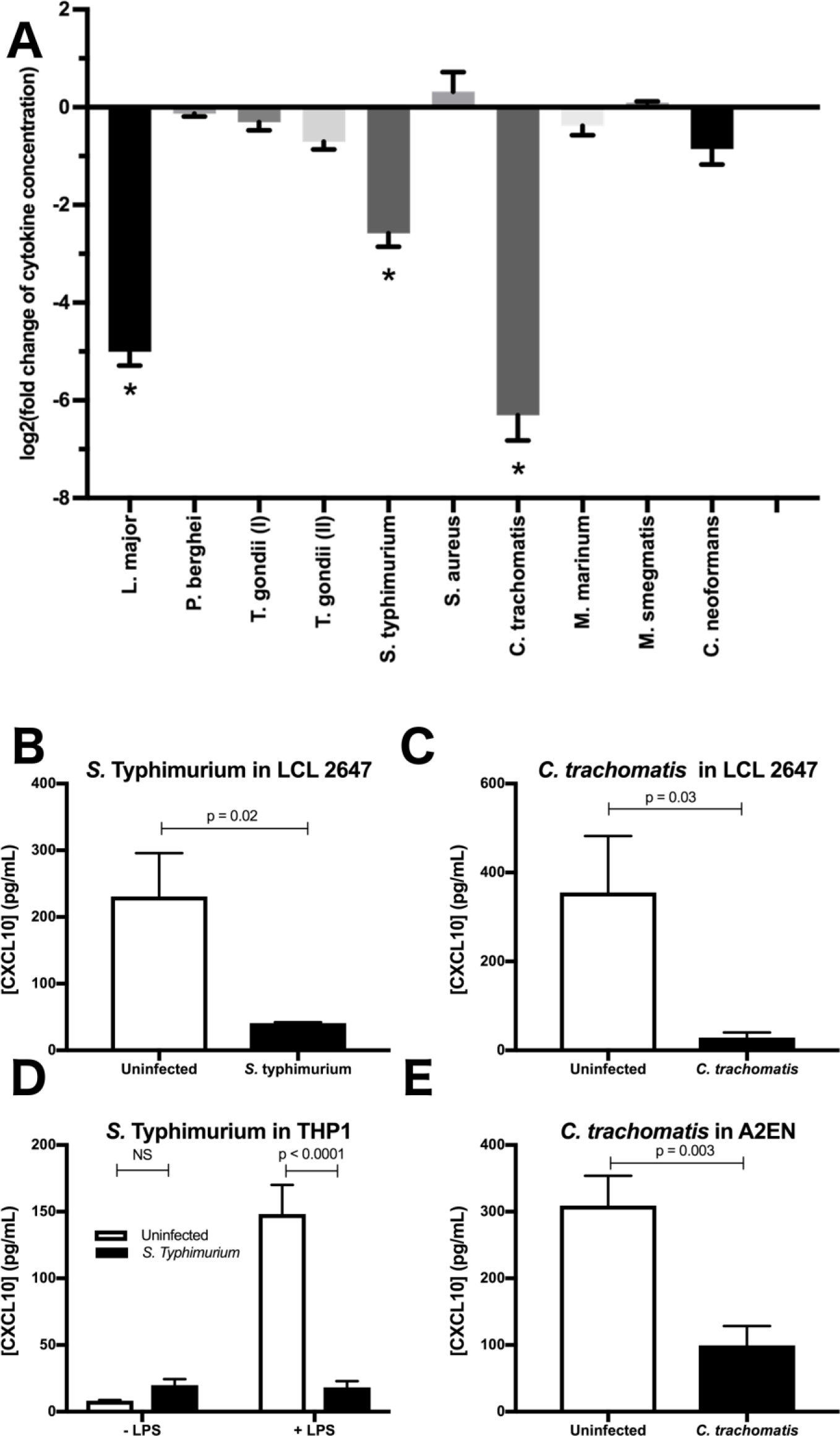
Multiple intracellular pathogens have evolved a mechanism for CXCL10 suppression. (A) LCL 18524 was used to screen *L. major* (p=0.0001), *P. berghei* (p=0.99), *T. gondii* I (RH) (p=0.44), *T. gondii* II (Prugniaud A7) (p=0.011), *S. enterica* serovar Typhimurium (p=0.0001), *S. aureus* (p=0.12), *C. trachomatis* (p=0.0001), *M. marinum* (p=0.37), *M. smegmatis* (p>0.99), *and C. neoformans* (p=0.010) for CXCL10 suppressing activity (n=2-4 for each pathogen). CXCL10 concentration was measured by ELISA and is represented as the log_2_ of fold change relative to uninfected controls. P values calculated by one-way ANOVA with Dunnett’s post-hoc test comparing non-log transformed values to 1, which would represent no change relative to uninfected. (*) represents p<0.01. (B-C) *S.* Typhimurium and *C. trachomatis* suppress CXCL10 in a second LCL. Infections were performed in the LCL HG02647 for *S.* Typhimurium (n=6; two experiments) and *C. trachomatis* (n=5; three experiments). Mean +/− standard error of the mean is plotted and P values calculated by Student’s *t-test*. (D) *S.* Typhimurium suppresses production of CXCL10 in THP-1 monocytes. THP-1 monocytes were stimulated with 1μg/mL of purified LPS from *S.* Typhimurium at the time of infection. CXCL10 concentration in culture supernatant at 24hpi was assayed by ELISA. Mean +/− standard error the mean is plotted, and P values calculated by two-way ANOVA with Tukey’s post-hoc test. (E) *C. trachomatis* suppresses CXCL10 in the human endocervical epithelial cell line A2EN. CXCL10 concentration in culture supernatant at 72hpi was assayed by ELISA. Mean +/− standard error of the mean is plotted and P values calculated by Student’s *t-test*.

Confirmation and characterization of CXCL10 suppression in different cell lines demonstrated that diverse intracellular pathogens impair chemokine accumulation. Using a second LCL, we confirmed that *S.* Typhimurium and *C. trachomatis* infection suppress CXCL10 (Fig. 5B-C). We then assessed the generalizability of CXCL10 suppression in host cell types known to be commonly infected by each pathogen. THP-1 monocytes stimulated with LPS upregulate significant production of CXCL10 protein, but infection with live *S.* Typhimurium dramatically impaired this CXCL10 induction (Fig. 5D). Similarly, the cervical epithelial cell line A2EN produces CXCL10 at baseline, but this is significantly reduced after infection with *C. trachomatis* (Fig. 5E). Thus, multiple intracellular pathogens that pose significant health burdens around the globe have independently evolved the ability to suppress CXCL10 in the cell types relevant to their infective niche.

## Discussion

We describe a mechanism used by intracellular pathogens to evade host chemokine response. Specifically, *L. major* can significantly reduce CXCL10 and impair its chemotactic activity through the matrix-metalloprotease, GP63. This strategy is likely to be highly beneficial to the parasite as CXCL10 protects against *L. major* (29), *L. amazonensis* (26), and *L. donovani* infection (27, 28). A similar phenotype of immune evasion that is shared by diverse intracellular pathogens points to a critical conserved role for CXCL10 in immunity to intracellular pathogens.

Consistent with CXCL10 playing a protective role during infection, multiple studies show that recruitment of CXCR3-expressing cells actively shapes the immune response. In response to *Leishmania* spp. specifically, CXCL10 is critical for the recruitment and activation of several cell types that contribute to the coordination of a protective type-1 immune response: natural killer (NK) cells, CD8+ T cells, dendritic cells, and CD4+ T_h_1 cells. With the early upregulation of *CXCL10* transcript (22), NK cells recruited during infection produce IFN_γ_ that contributes to T_h_1 differentiation (29, 56). Specific subsets of effector CD8+ T cells are recruited by CXCL10 after infection (23, 57). Finally, dendritic cells exposed to CXCL10 produce increased IL12, a cytokine that promotes T_h_1 polarization, and T_h_1 cells exposed to CXCL10 produce greater amounts of IFN_γ_ (58), a signal which infected macrophages require for efficient parasite killing (6). Beyond *Leishmania*, CXCR3-expressing cells have also been reported to play important roles in other infectious and inflammatory models (19, 23, 59–62). After infection with lymphocytic choriomeningitis virus, CXCR3 deletion leads to impaired production and localization of effector CD8+ T cells (63), and CXCL10 precisely coordinates effector CD8+ T cells to the site of *Toxoplasma gondii*, another intracellular eukaryotic pathogen (64). In response to the bacterial pathogen *S.* Typhimurium, which we identified as also suppressing CXCL10, mice have a significant expansion of CXCR3+ T_h_1 cells which border bacteria-rich granulomas in the spleen (43). These diverse examples highlight the importance of evading the CXCL10-CXCR3 signaling axis for pathogens.

Current limitations of parasite genetics as they relate to the complexity of GP63 related proteases may contribute to an incomplete picture of the impact of GP63 on chemokine suppression. First, sequencing *L. major* revealed proteins distantly homologous to GP63 (approximately 35% amino acid identity) on chromosomes 24 and 31, in addition to the tandem array of *gp63* genes on chromosome 10 (47, 65). These related proteases may suppress CXCL10 during different stages of infection or cleave an additional set of host substrates, even though they are not required for CXCL10 cleavage under our *in vitro* conditions. Second, *L. major Δgp63*+*1* has one of the seven chromosome-10 *gp63* copies maintained on a plasmid under G418 selection and optimized for expression in promastigotes (55), making the currently available GP63 strains sub-optimal for *in vivo* experiments. Despite these limitations, we demonstrate that GP63 cleavage of CXCL10 is selective, rapid, and renders the chemokine non-functional. Further investigation beyond the scope of this manuscript will be required to elucidate the implications of CXCL10 cleavage in other infection contexts and animal models.

An effective vaccine to protect against leishmaniasis has been a tantalizing strategy for disease control with unrealized potential due to an incomplete understanding of how the parasites interact with the immune system. Historically, inoculation with live parasites in unexposed areas of skin has effectively prevented future infections (66); however, this strategy poses significant risks (67–69) and subsequent vaccine development efforts failed to confer long-term protection in human studies (66). Recent studies highlight the importance of chemokine recruitment in mounting an efficient secondary immune response. Specifically, transcription of *Cxcl10* is upregulated in T resident-memory (T_rm_) cells after secondary infection, and antibody blockade of CXCR3 prevents recruitment of circulating CD4+ T cells to the site of infection (70–72). Together with our finding that CXCR3 substrates are cleaved by *L.* major, this suggests that one of the goals of vaccine development should be to overcome parasite-encoded CXCR3 escape upon secondary infection. Promisingly, GP63-specific CD4+ T cells elicit strong IFN_γ_ and T_h_1 responses (73) while GP63 based vaccines elicit long term immunity in mice that is correlated with T_h_1 responses (74–77); a phenotype that could be enhanced by anti-GP63 antibodies functionally blocking cleavage of CXCR3 ligands. A complete understanding of how the parasite alters chemokine recruitment upon secondary infection may facilitate development of a vaccine that can provide long term immunity to leishmaniasis.

The relevance of these insights into immune evasion is made more impactful by the observation that CXCL10 suppression has arisen in multiple intracellular pathogens. We found that *L. major, S.* Typhimurium, and *C. trachomatis* independently evolved the ability to suppress CXCL10, which indicates that suppression of CXCR3 inflammatory signaling is advantageous for multiple intracellular pathogens. In addition to *S.* Typhimurium and *C. trachomatis*, several other commensal and pathogenic bacteria have been reported to suppress CXCL10, including *Lactobacillus paracasei*, *Streptococcus pyogenes, Finegoldia magna*, and *Porphymonas gingivalis* (78–80). Similarly, the fungal pathogen *Candida albicans* produces a signaling molecule to inhibit *CXCL10* transcription (81). Among viruses, Hepatitis C virus (HCV) upregulates host proteases to modify CXCL10 (82), Epstein-Barr virus (EBV) decreases transcription through chromatin remodeling at the *CXCL10* locus (83), and Zika virus (ZIKV) blocks translation of CXCL10 (84, 85). The repeated and independent evolution of CXCL10 evasion suggests that this chemokine poses a significant evolutionary pressure on common human pathogens. These diverse pathogens heavily impact global morbidity and mortality. Understanding how pathogens manipulate the CXCR3 signaling axis to their advantage may enable therapeutic countermeasures that circumvent or prevent pathogen suppression of CXCR3 signaling.

## Materials/Methods

### Cell Lines

LCLs from the International HapMap Project (86) (GM18524 from Han Chinese in Beijing, China, GM19203 from Yoruba in Ibadan, Nigeria, GM7357 from Utah residents with Northern and Western European ancestry from the CEPH collection, and HG02647 of Gambian ancestry isolated in Gambia) were purchased from the Coriell Institute. LCLs were maintained at 37°C in a 5% CO2 atmosphere and were grown in RPMI 1640 media (Invitrogen) supplemented with 10% fetal bovine serum (FBS), 2 mM glutamine, 100 U/ml penicillin-G, and 100 mg/ml 790 streptomycin. THP-1 monocytes, originally from ATCC, were obtained from the Duke Cell Culture Facility and maintained in RPMI 1640 as described above. HEK293T cells were obtained from ATCC and maintained in DMEM complete media (Invitrogen) supplemented with 10% FBS, 100U/ml penicillin-G, and 100mg/ml 790 streptomycin. Jurkat cells (an immortalized T cell line) stably expressing CXCR3 were generated by transfecting a linearized pcDNA3.1 expression vector encoding *CXCR3* and resistance to Geneticin (G-418), selecting for transfected cells with 1000 μg/mL Geneticin, and collecting highly expressing CXCR3 cells by FACS. Cells were maintained in RPMI 1640 media (Sigma) supplemented with 10% FBS, 1% Penicillin/Streptomycin, 0.23% Glucose, 10mM HEPES, 1mM Sodium Pyruvate, and 250 μg/mL Geneticin. The A2EN cell line was provided by Raphael Valdivia and maintained in Keratinocyte serum free media (Gibco; 17005-042) supplemented with 10% heat-inactivated FBS, Epidermal Growth Factor 1-53, and Bovine Pituitary Extract.

### Pathogen culture and infections

*Leishmania* spp. were obtained from BEI (*L. major WT* ((MHOM/SN/74/Seidman), NR-48819), *L. major Δgp63* ((MHOM/SN/74/SD) *Δgp63 1-7*, NR-42489), *L. major Δgp63*+*1* (MHOM/SN/74/SD) *Δgp63 1-7* + *gp63-1*, NR-42490)). Parasites were maintained at 27°C in M199 media (Sigma-Aldrich, M1963), supplemented with 100u/ml penicillin/streptomycin, and 0.05% Hemin (Sigma-Aldrich, 51280). Cultures were split 1:20 every 5 days into 10mL of fresh culture media. To prepare parasites for infection, 8mL of a 5-day-old culture was spun at 1200g for 10 min and washed with 5mL of HBSS prior to counting promastigotes with a hemocytometer and resuspending at the indicated concentration. As relevant controls in using these *L. major* strains, we found the strains contained similar levels of metacyclic parasites based on flow cytometric measurement (87) and metacyclic enrichment with peanut agglutinin (Fig. S1).

For *Leishmania major* infections of LCLs and THP-1 monocytes, 1×10^5^ cells were placed in 100μl of RPMI 1640 assay media as described above, with no penicillin/streptomycin added. In the case of THP-1 monocytes, cells were then stimulated with 1μg/mL of LPS derived from *Salmonella enterica* serovar Typhimurium S-form` (Enzo Bioscience, ALX-581-011-L001). Finally, 1×10^6^ *L. major* promastigotes were added in 50μL of RPMI 1640 assay media for a multiplicity of infection (MOI) of 10. Culture supernatants and cell pellets were collected after 24 hours of infection. For phorbol 12-myristate 13-acetate (PMA) differentiation of THP-1 monocytes, 1.2×10^6^ cells were placed in 2mL of complete RPMI 1640 media supplemented with 100ng/mL of PMA for 8 hours after which the RPMI media was replaced and cells allowed to rest for 36 hours. Parasites were then washed and counted as described above and added at an MOI of 20. At 24 hours post-infection, the culture supernatant was removed, spun at 1200g for 10 minutes to separate extracellular parasites, and stored at −20C for downstream cytokine analysis. Cells were then washed 3 times with 1mL of PBS followed by one additional wash with 2mL of RPMI media to remove the remaining extracellular promastigotes. At 48 hours post infection the culture supernatant was collected and stored for downstream analysis. All cells were stored in 1mL of RNAprotect (Qiagen, 76526) at −20C for downstream RNA extraction (RNeasy Mini Kit, Qiagen, 74106) and qPCR analysis.

Screening GM18524 CXCL10 after infection with *Salmonella enterica* serovar Typhimurium 14028s, *Chlamydia trachomatis* serovar L2, and *Toxoplasma gondii* (RH and Prugniaud A7) were performed as described previously (88). For *Staphylococcus aureus*, LCLs were plated at 40,000 cells per 100μl RPMI assay media in 96-well plates prior to inoculation at an MOI of 10 with *S. aureus* Sanger-476. Cells were spun at 200xg for 5 minutes prior to incubation at 37°C for 1 hour. Gentamicin was added at 50µg/ml and then supernatant was collected at 24 hours. For *Cryptococcus neoformans*, LCLs were plated at 40,000 cells per 100μl RPMI assay media in 96-well plates prior to inoculation at an MOI of 5 with *C. neoformans* H99 strain. Cells were incubated at 37°C for 24 hours prior to collection of supernatant. For *Plasmodium berghei* infections, LCLs were plated at 40,000 cells per 100μl RPMI assay media in 96-well plates prior to inoculation with 17,000 *P. berghei*-Luciferase sporozoites isolated from *Anopheles stephensi* from the New York University Insectary Core Facility. Cells were spun at 1000×g for 10 minutes prior to incubation at 37°C for 48 hours. Cell death was monitored by 7AAD staining and quantified using a Guava easyCyte HT flow cytometer. To harvest supernatants, LCLs were centrifuged at 200xg for 5 minutes prior to removing supernatant and storing at −80°C prior to quantifying chemokines production by ELISA. For *Mycobacterium marinum* and *Mycobacterium smegmatis* infections, LCLs were plated at 40,000 cells per 100μl RPMI assay media without FBS and supplemented with 0.03% bovine serum albumin (BSA) prior to infection with 400,000 bacteria per well. Cells were spun at 100xg for 5 minutes prior to incubation at 33°C for 3 hours, after which 50µl of streptomycin in RPMI media was added for a final concentration of 200μg/ml streptomycin with 10% FBS, and incubation was continued at 33°C for 24 hours. Cell death was monitored by 7AAD staining and quantified using a Guava easyCyte HT flow cytometer. To harvest supernatants, LCLs were centrifuged at 200xg for 5 minutes prior to removing supernatant and storing at −80°C prior to quantifying chemokines by ELISA.

Confirmation of suppression by *S.* Typhimurium and *C. trachomatis* in LCL HG02647 was performed in 24 well plate format. For *S.* Typhimurium infection, 5×10^5^ cells were washed with antibiotic free RPMI assay media and plated in 500μl of RPMI assay media prior to infection at MOI 30. At 1 hour post infection, gentamycin was added at 50μg/mL to kill the remaining extracellular bacteria. At 2 hours post infection, gentamycin was diluted to 18μg/mL to prevent killing of intracellular bacteria. For *C. trachomatis* infection, 2×10^5^ cells were washed and plated in 500μl of RPMI assay media prior to infection at MOI 5 followed by centrifugation at 1500g for 30 minutes. For *S.* Typhimurium infection of THP-1 monocytes, cells were washed once with antibiotic free RPMI assay media and resuspended at a concentration of 1×10^5^ in 100μl of RPMI assay media on a 96-well plate. Cells were then treated with 1μg/mL of LPS diluted in RPMI assay media or the equivalent volume of media and *S.* typhimurium added at an MOI of 10. At 1 hour post infection, gentamycin was added at 50μg/mL. At 2 hours post infection, gentamycin was diluted to 25 μg/mL. For *C. trachomatis* infection of A2EN cells, 1×10^5^ cells were plated in a 96 well plate the day prior to infection. *C. trachomatis* was added at an MOI of 5 and centrifuged for 30 minutes at 1500g. For all *S.* Typhimurium infections, culture supernatants were harvested at 24 hours post infection. For *C. trachomatis* infection, culture supernatants were collected cells at 72 hours post infection to assess cytokine production.

### In vitro *T cell migration*

RPMI 1640 media (Sigma) was supplemented with 2% FBS and CXCL10 at a starting concentration of 100nM. This was pre-incubated at a ratio of 1:1 with conditioned media from either *L. major WT, Δgp63*, or *Δgp63*+*1*. After 12 hours of pre-incubation, 600µl of CXCL10/conditioned media mix was added to a 24 well plate. 500,000 Jurkat T cells stably transfected with *CXCR3* were seeded onto the apical membrane of the 5μm transwell insert (Corning, 3421), and allowed to incubate at 37°C for 4 hours. The transwell insert was removed and the concentration of cells in the basal chamber determined using a Guava easyCyte HT flow cytometer (Millipore).

### Expression of recombinant GP63 and site directed mutagenesis

Expression of CXCL10 and GP63 were performed by transfection in HEK293T cells. HEK293Ts were maintained in complete DMEM media supplemented with 10% FBS. Two days prior to transfection, 250,000 cells were washed and plated in a 6-well tissue culture treated plate in 2mL of serum free, FreeStyle 293 Expression Media (ThermoFisher, 12338018). One hour prior to transfection, media was replaced with fresh FreeStyle media. Transfection was performed with 2.5 total μg of endotoxin free plasmid DNA using the Lipofectamine 3000 Transfection Reagent Kit per manufacturer’s instructions. Transfected HEK293Ts were incubated at 37°C for 48 hours prior to harvesting culture supernatant and storing in polypropylene, low-binding tubes (Corning, 29442-578) at −80°C until use.

The CXCL10 plasmid was obtained from Origene (NM_001565), and contains C-terminal Myc and Flag epitope tags. For GP63, a codon optimized plasmid was obtained from OriGene on the pcDNA3.1/Hygro plasmid backbone. Following a kozak sequence and secrecon to enhance secretion (89–91), GP63-1 based on the *L. major* Fd sequence (Q4QHH2-1) was inserted with the *Leishmania* specific secretion signal and GPI anchor motif removed (V100-N577) (92), and epitope tagged with Myc and His sequences placed at the C-terminus. Point mutations in CXCL10 and GP63 were made using the Agilent QuikChange Site Directed Mutagenesis kit according to manufacturer’s instructions.

### Mass spectrometry

CXCL10 exposed to GP63 for 5 hours, along with a negative (untreated) control was delivered in PAGE loading buffer at an approximate concentration of 30 ng/uL. Mass spectrometry was carried out by the Duke Proteomics and Metabolomics Shared Resource. Molecular weight analysis of intact and cleaved CXCL10 from gel loading buffer was performed using a ZipChip CE system (908 Devices, Inc) coupled to a Q Exactive HF Orbitrap mass spectrometer (Thermo Scientific). Ammonium acetate was added to the sample to a final concentration 0.1 M, and 5µL of the loading buffer was pipetted manually into a HR ZipChip. Capillary electrophoresis (CE) separation was performed at 500 V/cm with a 30 second injection in Metabolite BGE (908 Devices, Inc). Mass spectrometry used positive electrospray with 120,000 Rs scan, 500-2000 m/z, 3e6 AGC target and 100 msec max ion injection time. Mass deconvolution was performed in Proteome Discoverer 2.2.

Tandem mass spectrometric sequencing of the cleaved and uncleaved fragments of CXCL10 after GP63 treatment, as well as an untreated control sample, were performed after gel separation on a 4-12% NuPAGE gel (Invitrogen). Gel bands were isolated after colloidal Coomassie staining, destained in acetonitrile/water, reduced with 10 mM DTT, alkylated with 20 mM iodoacetamide, and digested overnight at 37°C with 300 ng sequencing grade trypsin (Promega) in 50 mM ammonium bicarbonate at pH 8. Peptides were extracted in 1% formic acid and dried on a speedvac, then resuspended in a total of 10 µL 97/2/1 v/v/v water/acetonitrile/TFA. 4 µL of each sample was injected for analysis by LC-MS/MS using a 90 minute, 5-30% MeCN LC gradient and a top 12 DDA MS/MS method with MS1 at 120k and MS2 at 15k resolution. The data files were searched on Mascot v 2.5 with the UniProt database (downloaded November 2017) and *Homo sapiens* taxonomy selected, semitryptic specificity, along with fixed modification carbamidomethyl (C) and variable modifications oxidated (M), and deamidated (NQ). The results of the database searches were compiled into Scaffold v4 for curation. Using the search results as a spectral library, Skyline v4.1 was used to extract peak intensities for peptides which looked to be a part of the cleavage region (residues 74-91) or non-cleaved region (residues 48-68), in order to more definitively localize the specific cleavage location (Figure 2E). Intensity was expressed as the peak area normalized to the protein region from residues 29-52, in order to control for protein abundance differences between the samples. The Skyline file has been made publicly available at Panoramaweb.org (https://goo.gl/4xsLsF).

### Statistical analysis

All statistical analysis was performed using GraphPad Prism. Unpaired Student’s *t-test*, one-way ANOVA, and two-way ANOVA with Tukey’s post-hoc test were used as appropriate where indicated. The number of biological replicates (N) are indicated in the figure legend for each experiment and defined as follows. For *in vitro* cell culture and protein assessment each well of cells or chemokine prior to experimental manipulation (such as infection with parasite or addition of chemokine and/or inhibitor) was treated as a unique biological replicate. When technical replicates, repeated use of the same biological sample in a readout assay, were used they are indicated in the figure legend text and averaged values were combined into the single biological replicate prior to calculating statistics.

## Acknowledgments

ALA, KDG, ET, KJP, BHS, and DCK were supported by Duke Molecular Genetics & Microbiology startup funds, Duke University Whitehead Scholarship, Butler Pioneer Award, and NIH U19AI084044. ET was supported by the Duke Summer Research Opportunity Program (SROP). JSS and SR were supported by NIH GM122798 and the Burroughs Wellcome Fund Career Award for Medical Scientists. RLR was supported by NIH AI119004. We thank Robyn Guo for assistance with immunoblotting, Sarah Rains for assistance with sample prep for CXCL10 cleavage site identification, and Jeffrey S. Bourgeois for thoughtful discussion about experimental design and analysis. We thank the Duke University School of Medicine for the use of the Proteomics and Metabolomics Shared Resource.

## Author contributions

All authors critically reviewed the manuscript and contributed input to the final submission. ALA, DCK, KDG, and RLR wrote the manuscript. ALA, DCK, RLR, JSS, SR, and JWT contributed to strategy and project planning. ALA, KDG, ET, KJP, BHS, JSS, JWT, RLR, and DCK carried out experiments and analysis.

---

**Figure S1.**
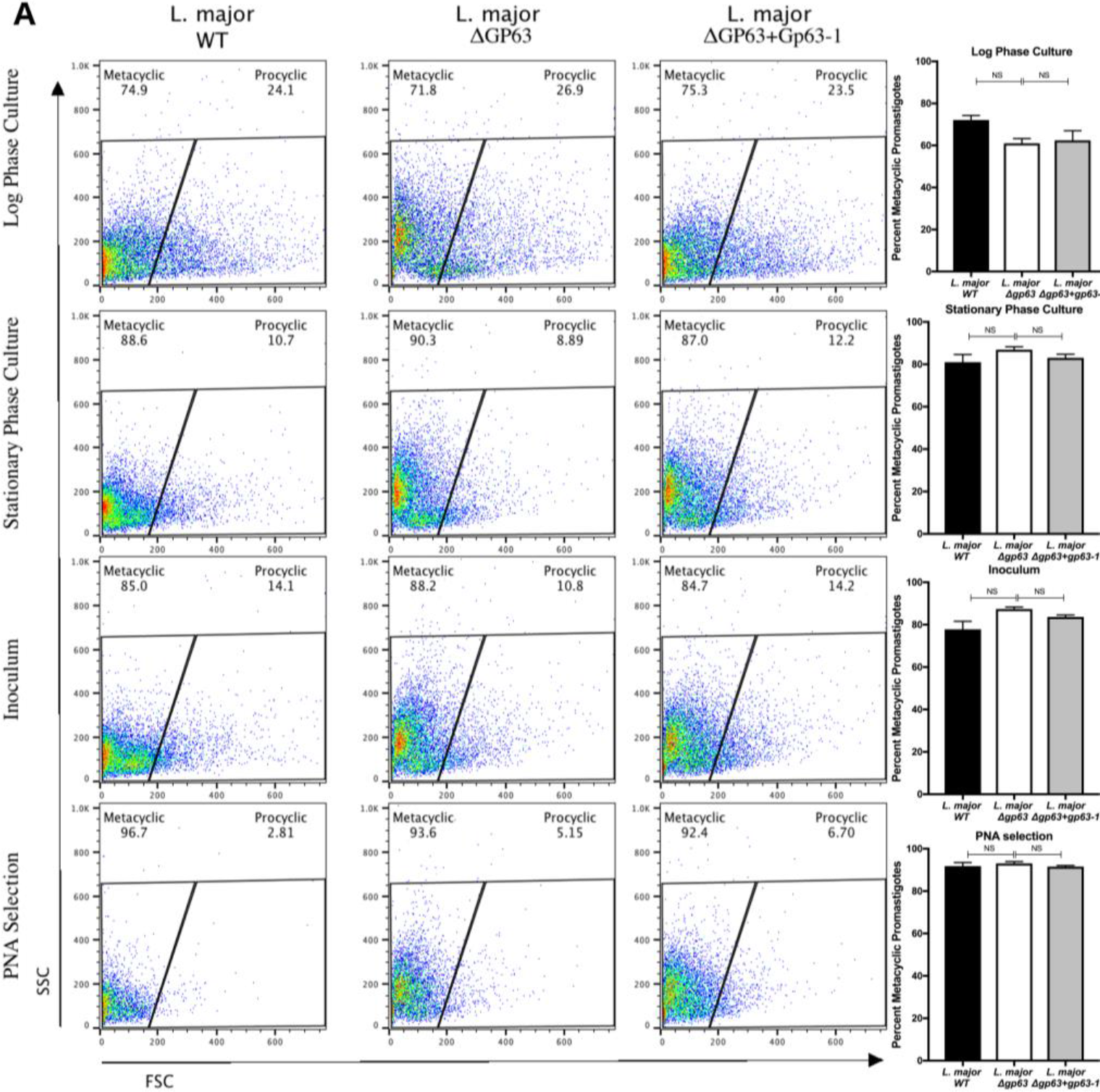
*L. major* WT, *L. major Δgp63*, and *L. major Δgp63*+*1* undergo comparable rates of metacyclogenesis. (A) Flow cytometry and selection with peanut agglutinin (PNA) demonstrate that the three strains of *L. major* used in this study did not have significantly different rates of metacyclogenesis. Parasites were analyzed using a Guava EasyCyte-HT flow cytometer and gated based on forward scatter (FSC) and side scatter (SSC) as previously described (87). Log-phase parasites were obtained from day 3 of culture, stationary phase parasites from day 5 of culture, the inoculum from day 5 culture after preparing parasites for infection as described in methods, and PNA selected from the inoculum after PNA selection. PNA selection was performed by incubating 1×10^8^ parasites in 100μg/mL of PNA (Vector Labs, L-1070-5) for 30 minutes at room temperature, followed by spinning for 5 minutes at 200xg, and taking the PNA-parasites in the supernatant for analysis. The PNA-parasites were then used as a control to define the gate for metacyclic promastigotes based on FSC and SSC. Conditions (n = 3 per group) were analyzed by one-way ANOVA with Tukeys post-hoc test. Not significant (NS) indicated p > 0.05.

## References

1. Reiner SL, Locksley RM. 1995. The regulation of immunity to Leishmania major. Annu Rev Immunol 13:151–177.

2. Alvar J, Velez ID, Bern C, Herrero M, Desjeux P, Cano J, Jannin J, den Boer M, Team WHOLC. 2012. Leishmaniasis worldwide and global estimates of its incidence. PLoS One 7:e35671.

3. Okwor I, Uzonna JE. 2009. Immunotherapy as a strategy for treatment of leishmaniasis: a review of the literature. Immunotherapy 1:765–776.

4. Hajishengallis G, Lambris JD. 2011. Microbial manipulation of receptor crosstalk in innate immunity. Nat Rev Immunol 11:187–200.

5. Finlay BB, McFadden G. 2006. Anti-immunology: evasion of the host immune system by bacterial and viral pathogens. Cell 124:767–782.

6. Scott P, Novais FO. 2016. Cutaneous leishmaniasis: immune responses in protection and pathogenesis. Nat Rev Immunol 16:581–592.

7. Kim CH, Rott L, Kunkel EJ, Genovese MC, Andrew DP, Wu L, Butcher EC. 2001. Rules of chemokine receptor association with T cell polarization in vivo. J Clin Invest 108:1331–1339.

8. Reiner SL, Locksley RM. 1993. Cytokines in the differentiation of Th1/Th2 CD4+ subsets in leishmaniasis. J Cell Biochem 53:323–328.

9. Heinzel FP, Sadick MD, Holaday BJ, Coffman RL, Locksley RM. 1989. Reciprocal expression of interferon gamma or interleukin 4 during the resolution or progression of murine leishmaniasis. Evidence for expansion of distinct helper T cell subsets. J Exp Med 169:59–72.

10. Scott P, Natovitz P, Coffman RL, Pearce E, Sher A. 1988. CD4+ T cell subsets in experimental cutaneous leishmaniasis. Mem Inst Oswaldo Cruz 83 Suppl 1:256–259.

11. Uzonna JE, Joyce KL, Scott P. 2004. Low dose Leishmania major promotes a transient T helper cell type 2 response that is down-regulated by interferon gamma-producing CD8+ T cells. J Exp Med 199:1559–1566.

12. Belkaid Y, Von Stebut E, Mendez S, Lira R, Caler E, Bertholet S, Udey MC, Sacks D. 2002. CD8+ T cells are required for primary immunity in C57BL/6 mice following low-dose, intradermal challenge with Leishmania major. J Immunol 168:3992–4000.

13. Muller I, Kropf P, Louis JA, Milon G. 1994. Expansion of gamma interferon-producing CD8+ T cells following secondary infection of mice immune to Leishmania major. Infect Immun 62:2575–2581.

14. Muller I, Kropf P, Etges RJ, Louis JA. 1993. Gamma interferon response in secondary Leishmania major infection: role of CD8+ T cells. Infect Immun 61:3730–3738.

15. Ritter U, Korner H. 2002. Divergent expression of inflammatory dermal chemokines in cutaneous leishmaniasis. Parasite Immunol 24:295–301.

16. Ajdary S, Alimohammadian MH, Eslami MB, Kemp K, Kharazmi A. 2000. Comparison of the immune profile of nonhealing cutaneous Leishmaniasis patients with those with active lesions and those who have recovered from infection. Infect Immun 68:1760–1764.

17. Carvalho EM, Correia Filho D, Bacellar O, Almeida RP, Lessa H, Rocha H. 1995. Characterization of the immune response in subjects with self-healing cutaneous leishmaniasis. Am J Trop Med Hyg 53:273–277.

18. Castellano LR, Filho DC, Argiro L, Dessein H, Prata A, Dessein A, Rodrigues V. 2009. Th1/Th2 immune responses are associated with active cutaneous leishmaniasis and clinical cure is associated with strong interferon-gamma production. Hum Immunol 70:383–390.

19. Groom JR, Luster AD. 2011. CXCR3 ligands: redundant, collaborative and antagonistic functions. Immunol Cell Biol 89:207–215.

20. Vargas-Inchaustegui DA, Hogg AE, Tulliano G, Llanos-Cuentas A, Arevalo J, Endsley JJ, Soong L. 2010. CXCL10 production by human monocytes in response to Leishmania braziliensis infection. Infect Immun 78:301–308.

21. Antoniazi S, Price HP, Kropf P, Freudenberg MA, Galanos C, Smith DF, Muller I. 2004. Chemokine gene expression in toll-like receptor-competent and -deficient mice infected with Leishmania major. Infect Immun 72:5168–5174.

22. Zaph C, Scott P. 2003. Interleukin-12 regulates chemokine gene expression during the early immune response to Leishmania major. Infect Immun 71:1587–1589.

23. Oghumu S, Dong R, Varikuti S, Shawler T, Kampfrath T, Terrazas CA, Lezama-Davila C, Ahmer BM, Whitacre CC, Rajagopalan S, Locksley R, Sharpe AH, Satoskar AR. 2013. Distinct populations of innate CD8+ T cells revealed in a CXCR3 reporter mouse. J Immunol 190:2229–2240.

24. Barbi J, Oghumu S, Rosas LE, Carlson T, Lu B, Gerard C, Lezama-Davila CM, Satoskar AR. 2007. Lack of CXCR3 delays the development of hepatic inflammation but does not impair resistance to Leishmania donovani. J Infect Dis 195:1713–1717.

25. Barbi J, Brombacher F, Satoskar AR. 2008. T cells from Leishmania major-susceptible BALB/c mice have a defect in efficiently up-regulating CXCR3 upon activation. J Immunol 181:4613–4620.

26. Vasquez RE, Soong L. 2006. CXCL10/gamma interferon-inducible protein 10-mediated protection against Leishmania amazonensis infection in mice. Infect Immun 74:6769–6777.

27. Gupta G, Bhattacharjee S, Bhattacharyya S, Bhattacharya P, Adhikari A, Mukherjee A, Bhattacharyya Majumdar S, Majumdar S. 2009. CXC chemokine-mediated protection against visceral leishmaniasis: involvement of the proinflammatory response. J Infect Dis 200:1300–1310.

28. Gupta G, Majumdar S, Adhikari A, Bhattacharya P, Mukherjee AK, Majumdar SB, Majumdar S. 2011. Treatment with IP-10 induces host-protective immune response by regulating the T regulatory cell functioning in Leishmania donovani-infected mice. Med Microbiol Immunol 200:241–253.

29. Vester B, Muller K, Solbach W, Laskay T. 1999. Early gene expression of NK cell-activating chemokines in mice resistant to Leishmania major. Infect Immun 67:3155–3159.

30. Gondek DC, Roan NR, Starnbach MN. 2009. T cell responses in the absence of IFN-gamma exacerbate uterine infection with Chlamydia trachomatis. J Immunol 183:1313–1319.

31. Morrison RP, Caldwell HD. 2002. Immunity to murine chlamydial genital infection. Infect Immun 70:2741–2751.

32. Morrison SG, Farris CM, Sturdevant GL, Whitmire WM, Morrison RP. 2011. Murine Chlamydia trachomatis genital infection is unaltered by depletion of CD4+ T cells and diminished adaptive immunity. J Infect Dis 203:1120–1128.

33. Perry LL, Feilzer K, Caldwell HD. 1997. Immunity to Chlamydia trachomatis is mediated by T helper 1 cells through IFN-gamma-dependent and -independent pathways. J Immunol 158:3344–3352.

34. Rank RG, Lacy HM, Goodwin A, Sikes J, Whittimore J, Wyrick PB, Nagarajan UM. 2010. Host chemokine and cytokine response in the endocervix within the first developmental cycle of Chlamydia muridarum. Infect Immun 78:536–544.

35. Lijek RS, Helble JD, Olive AJ, Seiger KW, Starnbach MN. 2018. Pathology after Chlamydia trachomatis infection is driven by nonprotective immune cells that are distinct from protective populations. Proc Natl Acad Sci U S A 115:2216–2221.

36. Maxion HK, Kelly KA. 2002. Chemokine expression patterns differ within anatomically distinct regions of the genital tract during Chlamydia trachomatis infection. Infect Immun 70:1538–1546.

37. Hess J, Ladel C, Miko D, Kaufmann SH. 1996. Salmonella typhimurium aroA-infection in gene-targeted immunodeficient mice: major role of CD4+ TCR-alpha beta cells and IFN-gamma in bacterial clearance independent of intracellular location. J Immunol 156:3321–3326.

38. Ravindran R, Foley J, Stoklasek T, Glimcher LH, McSorley SJ. 2005. Expression of T-bet by CD4 T cells is essential for resistance to Salmonella infection. J Immunol 175:4603–4610.

39. Gilchrist JJ, MacLennan CA, Hill AV. 2015. Genetic susceptibility to invasive Salmonella disease. Nat Rev Immunol 15:452–463.

40. Saliba AE, Li L, Westermann AJ, Appenzeller S, Stapels DA, Schulte LN, Helaine S, Vogel J. 2016. Single-cell RNA-seq ties macrophage polarization to growth rate of intracellular Salmonella. Nat Microbiol 2:16206.

41. Jul 2015. 7. Replication of Salmonella enterica Serovar Typhimurium in Human Monocyte-Derived Macrophages. Infect Immun, 83.2661–2671. http://iai.asm.org/lookup/doi/10.1128/IAI.00033-15.

42. Martinez FO, Gordon S, Locati M, Mantovani A. 2006. Transcriptional profiling of the human monocyte-to-macrophage differentiation and polarization: new molecules and patterns of gene expression. J Immunol 177:7303–7311.

43. Goldberg MF, Roeske EK, Ward LN, Pengo T, Dileepan T, Kotov DI, Jenkins MK. 2018. Salmonella Persist in Activated Macrophages in T Cell-Sparse Granulomas but Are Contained by Surrounding CXCR3 Ligand-Positioned Th1 Cells. Immunity 49:1090–1102 e1097.

44. Chami B, Yeung A, Buckland M, Liu H, G MF, Tao K, Bao S. 2017. CXCR3 plays a critical role for host protection against Salmonellosis. Sci Rep 7:10181.

45. Khan IA, MacLean JA, Lee FS, Casciotti L, DeHaan E, Schwartzman JD, Luster AD. 2000. IP-10 is critical for effector T cell trafficking and host survival in Toxoplasma gondii infection. Immunity 12:483–494.

46. Olive AJ, Gondek DC, Starnbach MN. 2011. CXCR3 and CCR5 are both required for T cell-mediated protection against C. trachomatis infection in the murine genital mucosa. Mucosal Immunol 4:208–216.

47. Valdivia HO, Scholte LL, Oliveira G, Gabaldon T, Bartholomeu DC. 2015. The Leishmania metaphylome: a comprehensive survey of Leishmania protein phylogenetic relationships. BMC Genomics 16:887.

48. Olivier M, Atayde VD, Isnard A, Hassani K, Shio MT. 2012. Leishmania virulence factors: focus on the metalloprotease GP63. Microbes Infect 14:1377–1389.

49. Voth BR, Kelly BL, Joshi PB, Ivens AC, McMaster WR. 1998. Differentially expressed Leishmania major gp63 genes encode cell surface leishmanolysin with distinct signals for glycosylphosphatidylinositol attachment. Mol Biochem Parasitol 93:31–41.

50. Fernandes MC, Dillon LA, Belew AT, Bravo HC, Mosser DM, El-Sayed NM. 2016. Dual Transcriptome Profiling of Leishmania-Infected Human Macrophages Reveals Distinct Reprogramming Signatures. MBio 7.

51. Chaudhuri G, Chaudhuri M, Pan A, Chang KP. 1989. Surface acid proteinase (gp63) of Leishmania mexicana. A metalloenzyme capable of protecting liposome-encapsulated proteins from phagolysosomal degradation by macrophages. J Biol Chem 264:7483–7489.

52. Joshi PB, Kelly BL, Kamhawi S, Sacks DL, McMaster WR. 2002. Targeted gene deletion in Leishmania major identifies leishmanolysin (GP63) as a virulence factor. Mol Biochem Parasitol 120:33–40.

53. Bouvier J, Schneider P, Etges R, Bordier C. 1990. Peptide substrate specificity of the membrane-bound metalloprotease of Leishmania. Biochemistry 29:10113–10119.

54. Booth V, Keizer DW, Kamphuis MB, Clark-Lewis I, Sykes BD. 2002. The CXCR3 binding chemokine IP-10/CXCL10: structure and receptor interactions. Biochemistry 41:10418–10425.

55. Joshi PB, Sacks DL, Modi G, McMaster WR. 1998. Targeted gene deletion of Leishmania major genes encoding developmental stage-specific leishmanolysin (GP63). Mol Microbiol 27:519–530.

56. Muller K, van Zandbergen G, Hansen B, Laufs H, Jahnke N, Solbach W, Laskay T. 2001. Chemokines, natural killer cells and granulocytes in the early course of Leishmania major infection in mice. Med Microbiol Immunol 190:73–76.

57. Majumder S, Bhattacharjee S, Paul Chowdhury B, Majumdar S. 2012. CXCL10 is critical for the generation of protective CD8 T cell response induced by antigen pulsed CpG-ODN activated dendritic cells. PLoS One 7:e48727.

58. Vasquez RE, Xin L, Soong L. 2008. Effects of CXCL10 on dendritic cell and CD4+ T-cell functions during Leishmania amazonensis infection. Infect Immun 76:161–169.

59. Qin S, Rottman JB, Myers P, Kassam N, Weinblatt M, Loetscher M, Koch AE, Moser B, Mackay CR. 1998. The chemokine receptors CXCR3 and CCR5 mark subsets of T cells associated with certain inflammatory reactions. J Clin Invest 101:746–754.

60. Thomas SY, Hou R, Boyson JE, Means TK, Hess C, Olson DP, Strominger JL, Brenner MB, Gumperz JE, Wilson SB, Luster AD. 2003. CD1d-restricted NKT cells express a chemokine receptor profile indicative of Th1-type inflammatory homing cells. J Immunol 171:2571–2580.

61. Cella M, Jarrossay D, Facchetti F, Alebardi O, Nakajima H, Lanzavecchia A, Colonna M. 1999. Plasmacytoid monocytes migrate to inflamed lymph nodes and produce large amounts of type I interferon. Nat Med 5:919–923.

62. Nanki T, Takada K, Komano Y, Morio T, Kanegane H, Nakajima A, Lipsky PE, Miyasaka N. 2009. Chemokine receptor expression and functional effects of chemokines on B cells: implication in the pathogenesis of rheumatoid arthritis. Arthritis Res Ther 11:R149.

63. Hu JK, Kagari T, Clingan JM, Matloubian M. 2011. Expression of chemokine receptor CXCR3 on T cells affects the balance between effector and memory CD8 T-cell generation. Proc Natl Acad Sci U S A 108:E118–127.

64. Harris TH, Banigan EJ, Christian DA, Konradt C, Wojno EDT, Norose K, Wilson EH, John B, Weninger W, Luster AD, Liu AJ, Hunter CA. 2012. Generalized Levy walks and the role of chemokines in migration of effector CD8(+) T cells. Nature 486:545–U145.

65. Ivens AC, Peacock CS, Worthey EA, Murphy L, Aggarwal G, Berriman M, Sisk E, Rajandream MA, Adlem E, Aert R, Anupama A, Apostolou Z, Attipoe P, Bason N, Bauser C, Beck A, Beverley SM, Bianchettin G, Borzym K, Bothe G, Bruschi CV, Collins M, Cadag E, Ciarloni L, Clayton C, Coulson RM, Cronin A, Cruz AK, Davies RM, De Gaudenzi J, Dobson DE, Duesterhoeft A, Fazelina G, Fosker N, Frasch AC, Fraser A, Fuchs M, Gabel C, Goble A, Goffeau A, Harris D, Hertz-Fowler C, Hilbert H, Horn D, Huang Y, Klages S, Knights A, Kube M, Larke N, Litvin L, et al. 2005. The genome of the kinetoplastid parasite, Leishmania major. Science 309:436–442.

66. Seyed N, Peters NC, Rafati S. 2018. Translating Observations From Leishmanization Into Non-Living Vaccines: The Potential of Dendritic Cell-Based Vaccination Strategies Against Leishmania. Front Immunol 9:1227.

67. Singh S. 2014. Changing trends in the epidemiology, clinical presentation, and diagnosis of Leishmania-HIV co-infection in India. Int J Infect Dis 29:103–112.

68. Lindoso JA, Cota GF, da Cruz AM, Goto H, Maia-Elkhoury AN, Romero GA, de Sousa-Gomes ML, Santos-Oliveira JR, Rabello A. 2014. Visceral leishmaniasis and HIV coinfection in Latin America. PLoS Negl Trop Dis 8:e3136.

69. Monge-Maillo B, Norman FF, Cruz I, Alvar J, Lopez-Velez R. 2014. Visceral leishmaniasis and HIV coinfection in the Mediterranean region. PLoS Negl Trop Dis 8:e3021.

70. Romano A, Carneiro MBH, Doria NA, Roma EH, Ribeiro-Gomes FL, Inbar E, Lee SH, Mendez J, Paun A, Sacks DL, Peters NC. 2017. Divergent roles for Ly6C+CCR2+CX3CR1+ inflammatory monocytes during primary or secondary infection of the skin with the intra-phagosomal pathogen Leishmania major. PLoS Pathog 13:e1006479.

71. Glennie ND, Scott P. 2016. Memory T cells in cutaneous leishmaniasis. Cell Immunol 309:50–54.

72. Glennie ND, Yeramilli VA, Beiting DP, Volk SW, Weaver CT, Scott P. 2015. Skin-resident memory CD4+ T cells enhance protection against Leishmania major infection. J Exp Med 212:1405–1414.

73. Julia V, Glaichenhaus N. 1999. CD4(+) T cells which react to the Leishmania major LACK antigen rapidly secrete interleukin-4 and are detrimental to the host in resistant B10.D2 mice. Infect Immun 67:3641–3644.

74. Mazumder S, Maji M, Das A, Ali N. 2011. Potency, efficacy and durability of DNA/DNA, DNA/protein and protein/protein based vaccination using gp63 against Leishmania donovani in BALB/c mice. PLoS One 6:e14644.

75. Mazumder S, Maji M, Ali N. 2011. Potentiating effects of MPL on DSPC bearing cationic liposomes promote recombinant GP63 vaccine efficacy: high immunogenicity and protection. PLoS Negl Trop Dis 5:e1429.

76. Bhowmick S, Ravindran R, Ali N. 2008. gp63 in stable cationic liposomes confers sustained vaccine Immunity to susceptible BALB/c mice infected with Leishmania donovani. Infect Immun 76:1003–1015.

77. Sachdeva R, Banerjea AC, Malla N, Dubey ML. 2009. Immunogenicity and efficacy of single antigen Gp63, polytope and polytopeHSP70 DNA vaccines against visceral Leishmaniasis in experimental mouse model. PLoS One 4:e7880.

78. von Schillde MA, Hormannsperger G, Weiher M, Alpert CA, Hahne H, Bauerl C, van Huynegem K, Steidler L, Hrncir T, Perez-Martinez G, Kuster B, Haller D. 2012. Lactocepin secreted by Lactobacillus exerts anti-inflammatory effects by selectively degrading proinflammatory chemokines. Cell Host Microbe 11:387–396.

79. Karlsson C, Eliasson M, Olin AI, Morgelin M, Karlsson A, Malmsten M, Egesten A, Frick IM. 2009. SufA of the opportunistic pathogen finegoldia magna modulates actions of the antibacterial chemokine MIG/CXCL9, promoting bacterial survival during epithelial inflammation. J Biol Chem 284:29499–29508.

80. Jauregui CE, Wang Q, Wright CJ, Takeuchi H, Uriarte SM, Lamont RJ. 2013. Suppression of T-cell chemokines by Porphyromonas gingivalis. Infect Immun 81:2288–2295.

81. Shiraki Y, Ishibashi Y, Hiruma M, Nishikawa A, Ikeda S. 2008. Candida albicans abrogates the expression of interferon-gamma-inducible protein-10 in human keratinocytes. FEMS Immunol Med Microbiol 54:122–128.

82. Casrouge A, Decalf J, Ahloulay M, Lababidi C, Mansour H, Vallet-Pichard A, Mallet V, Mottez E, Mapes J, Fontanet A, Pol S, Albert ML. 2011. Evidence for an antagonist form of the chemokine CXCL10 in patients chronically infected with HCV. J Clin Invest 121:308–317.

83. Harth-Hertle ML, Scholz BA, Erhard F, Glaser LV, Dolken L, Zimmer R, Kempkes B. 2013. Inactivation of intergenic enhancers by EBNA3A initiates and maintains polycomb signatures across a chromatin domain encoding CXCL10 and CXCL9. PLoS Pathog 9:e1003638.

84. Bowen JR, Quicke KM, Maddur MS, O'Neal JT, McDonald CE, Fedorova NB, Puri V, Shabman RS, Pulendran B, Suthar MS. 2017. Zika Virus Antagonizes Type I Interferon Responses during Infection of Human Dendritic Cells. PLoS Pathog 13:e1006164.

85. Chaudhary V, Yuen KS, Chan JF, Chan CP, Wang PH, Cai JP, Zhang S, Liang M, Kok KH, Chan CP, Yuen KY, Jin DY. 2017. Selective Activation of Type II Interferon Signaling by Zika Virus NS5 Protein. J Virol 91.

86. Consortium IH. 2005. A haplotype map of the human genome. Nature 437:1299–1320.

87. Saraiva EM, Pinto-da-Silva LH, Wanderley JL, Bonomo AC, Barcinski MA, Moreira ME. 2005. Flow cytometric assessment of Leishmania spp metacyclic differentiation: validation by morphological features and specific markers. Exp Parasitol 110:39–47.

88. Wang L, Pittman KJ, Barker JR, Salinas RE, Stanaway IB, Williams GD, Carroll RJ, Balmat T, Ingham A, Gopalakrishnan AM, Gibbs KD, Antonia AL, e MN, Heitman J, Lee SC, Jarvik GP, Denny JC, Horner SM, DeLong MR, Valdivia RH, Crosslin DR, Ko DC. 2018. An Atlas of Genetic Variation Linking Pathogen-Induced Cellular Traits to Human Disease. Cell Host Microbe 24:308–323 e306.

89. Kozak M. 1989. The scanning model for translation: an update. J Cell Biol 108:229–241.

90. Barash S, Wang W, Shi Y. 2002. Human secretory signal peptide description by hidden Markov model and generation of a strong artificial signal peptide for secreted protein expression. Biochem Biophys Res Commun 294:835–842.

91. Guler-Gane G, Kidd S, Sridharan S, Vaughan TJ, Wilkinson TC, Tigue NJ. 2016. Overcoming the Refractory Expression of Secreted Recombinant Proteins in Mammalian Cells through Modification of the Signal Peptide and Adjacent Amino Acids. PLoS One 11:e0155340.

92. Schlagenhauf E, Etges R, Metcalf P. 1998. The crystal structure of the Leishmania major surface proteinase leishmanolysin (gp63). Structure 6:1035–1046.

